# Extreme nuclear branching in healthy epidermal cells of the *Xenopus* tail fin

**DOI:** 10.1101/364471

**Authors:** Hannah E. Arbach, Marcus Harland-Dunaway, Jessica K. Chang, Andrea E. Wills

## Abstract

Changes in nuclear morphology contribute to regulation of complex cell properties, including differentiation and tissue elasticity. Perturbations of nuclear morphology are associated with pathologies that include, progeria, cancer, and muscular dystrophy. The mechanisms governing nuclear shape changes in healthy cells remain poorly understood, partially because there are few healthy models of nuclear shape variation. Here, we introduce nuclear branching in epidermal fin cells of *Xenopus tropicalis* as a model for extreme variation of nuclear morphology in a diverse population of healthy cells. We find that nuclear branching arises and elaborates during embryonic development. They contain broadly distributed marks of transcriptionally active chromatin and heterochromatin and have active cell cycles. We find that nuclear branches are disrupted by loss of filamentous actin and depend on epidermal expression of the nuclear lamina protein Lamin B1. Inhibition of nuclear branching disrupts fin morphology, suggesting that nuclear branching may be involved in fin development. This study introduces the nuclei of the fin as a powerful new model for extreme nuclear morphology in healthy cells to complement studies of nuclear shape variation in pathological contexts.

**List of abbreviations and symbols:** LINC-Linker of Nucleoskeleton and Cytoskeleton, HGPS – Hutchinson-Gilford Progeria Syndrome, TEM – Transmission electron microscopy, PH3 – Phosphorylated Histone 3, Lat B – Latrunculin B, WT-Wild type, Cyto D-Cytochalasin D, *Lmnb1* CRISPR – *lmnb1* mutants generated by CRISPR/Cas9, E-Lmnb1 CRISPR – epidermal specific *lmnb1* mutants generated by CRISPR/Cas9, Scrmbl-tadpoles injected with a scrambled version of the *lmnb1* targeted sgRNA, Lmnb1-rod – dominant-negative Lamin B1 containing only the rod domain

**Summary Statement:** Nuclei are highly branched throughout the heterogeneous population of healthy epidermal cells that comprise the *Xenopus* tail fin periphery, and disruption of nuclear branching mechanisms results in improper fin morphology.

## Introduction

Nuclear shape is highly conserved across cell types and species. Most healthy cells have round or ellipsoid nuclei. A few healthy cell types exhibit non-ellipsoid morphologies, including neutrophils, which have a distinct lobular structure that allows them to extravasate to areas with damaged tissue (Pillay et al., 2013; Rowat et al., 2013). Frequently, perturbations in nuclear morphology are associated with disease. Well-studied examples include progeria (Chen et al., 2014; Dahl et al., 2006; Goldman et al., 2004; Schirmer et al., 2001; Verstraeten et al., 2008), muscular dystrophy (Bonne et al., 1999), neurodegeneration (Frost et al., 2016), and cancers (Denais and Lammerding, 2014; Fu et al., 2012; Shah et al., 2013). HeLa cells in particular are a model for nuclear morphological variation, which includes blebbing and ruffling of the nuclear membrane and dysregulation of multiple nucleoskeletal components (Wiggan et al., 2017). However, it is largely unclear what general mechanisms allow cells to acquire non-ellipsoid nuclear morphologies or how these morphologies could influence tissue and cellular function. One barrier to understanding extreme morphological variation of the nucleus is the dearth of models where nuclear morphology varies in the absence of disease. Here we characterize epidermal cells in the fin margin of *Xenopus tropicalis* tadpoles that have a non-ellipsoid, branched nuclear architecture. These striking nuclear morphologies arise during tail development and persist late into metamorphosis.

The nucleus derives its shape from interactions between the nucleoskeleton and the actin cytoskeleton. The nucleoskeleton is a complex network of Lamin filaments, associated proteins, and the LINC (**LI**nker of **N**ucleoskeleton and **C**ytoskeleton) complex (Chang et al., 2015; Chen et al., 2014; Davidson and Lammerding, 2014; Denais and Lammerding, 2014; Fu et al., 2012; Goldman et al., 2004; Schirmer et al., 2001; Vergnes et al., 2004; Zwerger et al., 2013). Alterations in nuclear lamina composition, particularly the relative levels of A-type and B-type Lamins, enable changes in not only nuclear shape but also nuclear deformability (Swift et al., 2013). Changes in this ratio allow the formation of nuclear lobes and a highly deformable nuclear envelope in neutrophils, which in turn enable passage through small capillaries. Perturbation of B-type Lamins or their receptors has deleterious effect on neutrophil migration (Dreesen et al., 2013; Rowat et al., 2013). More recent studies of interactions between perinuclear actin and the nuclear envelope have also clarified that the rigidity of the actin cap and degree of actin polymerization directly affect nuclear shape and tissue stiffness (Swift et al., 2013; Wiggan et al., 2017). Variation in nuclear morphology is therefore predicted to have consequences for the biophysical function of the associated tissue, although relatively little is known about the mechanism by which other nuclear functions are modulated or constrained by extreme shape change (Dahl et al., 2006; Pajerowski et al., 2007; Rowat et al., 2013; Zwerger et al., 2013).

The structural organization of the nuclear lamina scaffolds functional domains within chromatin and serves to protect the genome (Peric-Hupkes et al., 2010; Shah et al., 2013; Solovei et al., 2013). Chromatin-lamina interactions are important for appropriate gene regulation. Canonically, heterochromatin or repressed regions of the genome are associated with the nuclear lamina (Fraser et al., 2015; Mattout et al., 2015a; Peric-Hupkes et al., 2010). Alterations in heterochromatin propagation is linked to changes in nuclear morphology caused by laminopathies (Davidson and Lammerding, 2014; Dreesen et al., 2013; Perovanovic et al., 2016; Shah et al., 2013). Hutchinson-Gilford Progeria Syndrome (HGPS) is a laminopathy that causes premature aging and is associated with mutations in LMNA that disrupt prelamin A cleavage, leading to gross changes in nuclear morphology (Goldman *et al*., 2004; Dahl *et al*., 2006; Verstraeten *et al*., 2008; Chen *et al*., 2014). As cultured cells with HGPS *lemma* mutations undergo more passages, they acquire progressively more nuclear ruffling and alterations of heterochromatin, resembling senescent cells rather than proliferative cells. Similar alterations in heterochromatic regions are seen in Lamin B1-depleted cells and cancer cells (Perovanovic et al., 2016; Shah et al., 2013). This suggests that alteration of the nucleoskeleton can contribute to large-scale changes in chromatin reorganization and gene expression that contribute to aging or other pathologies.

In this study we have characterized nuclear branching in the fin epithelium of *Xenopus* tadpoles. The thin epithelium of the tadpole is made up of flattened epidermal cells that overlie a mesenchymal core (Tucker and Slack, 2004). Its specialized cell biological and biophysical properties allow rapid regeneration and sinusoidal swimming movements. We show that branched morphologies of the nuclear lumen, chromatin, and nuclear lamina arises during development in a heterogenous population of epidermal cells that make up the fin periphery. Cells with branched nuclei contain epigenetic marks of active enhancers and inactive chromatin throughout the nucleoplasm, additionally these cells have active cell cycles. We find that actin filaments, but not polymerized microtubules are necessary to maintain branched nuclear morphology. We also find that functional epidermal Lamin B1 is required for both nuclear branching and for proper development of the fin and tail.

## Results

### Nuclei in the tail of *Xenopus tropicalis* are branched

Although *Xenopus* has long served as a model for epidermal cell biology and nuclear composition, there has been little examination of nuclear morphology in the differentiated fin. We conducted whole-embryo DAPI stains of the *Xenopus tropicalis* tadpole fin, which revealed an unexpected elaborately branched distribution of DNA in the fin marginal cells (Fig. 1A). Although examples of potentially branched nuclear morphologies can be observed in earlier literature, these have not been described in detail (Davis and Kirschner, 2000). Our first goal was to establish whether branching was confined to chromatin or was shared by the nuclear lumen and envelope (Fig. 1B). To this end, cleavage-stage embryos were injected with either a cocktail of *h2b-rfp* and *lmnb3-gfp* mRNA to label histones (chromatin) and the nuclear lamina respectively, or with GFP bearing a nuclear localization signal (Nuclear GFP) (Fig. 1C). Tadpoles were reared to NF stage 41 and then live images were taken in the anterior-most third of the fin margin. Nuclear GFP confirmed that the nuclear lumen in cells of the fin margin is also highly branched structure (Fig. 1D). Consistent with our observations for DAPI, we find that H2B-RFP has a branched distribution in the nuclei of fin margin cells. Lamin B3-GFP localization showed that the nuclear compartment is also branched (Fig. 1D). Thus, the entire nucleus of fin marginal cells is branched, including the lamina, lumen, and chromatin.

**Figure 1.**
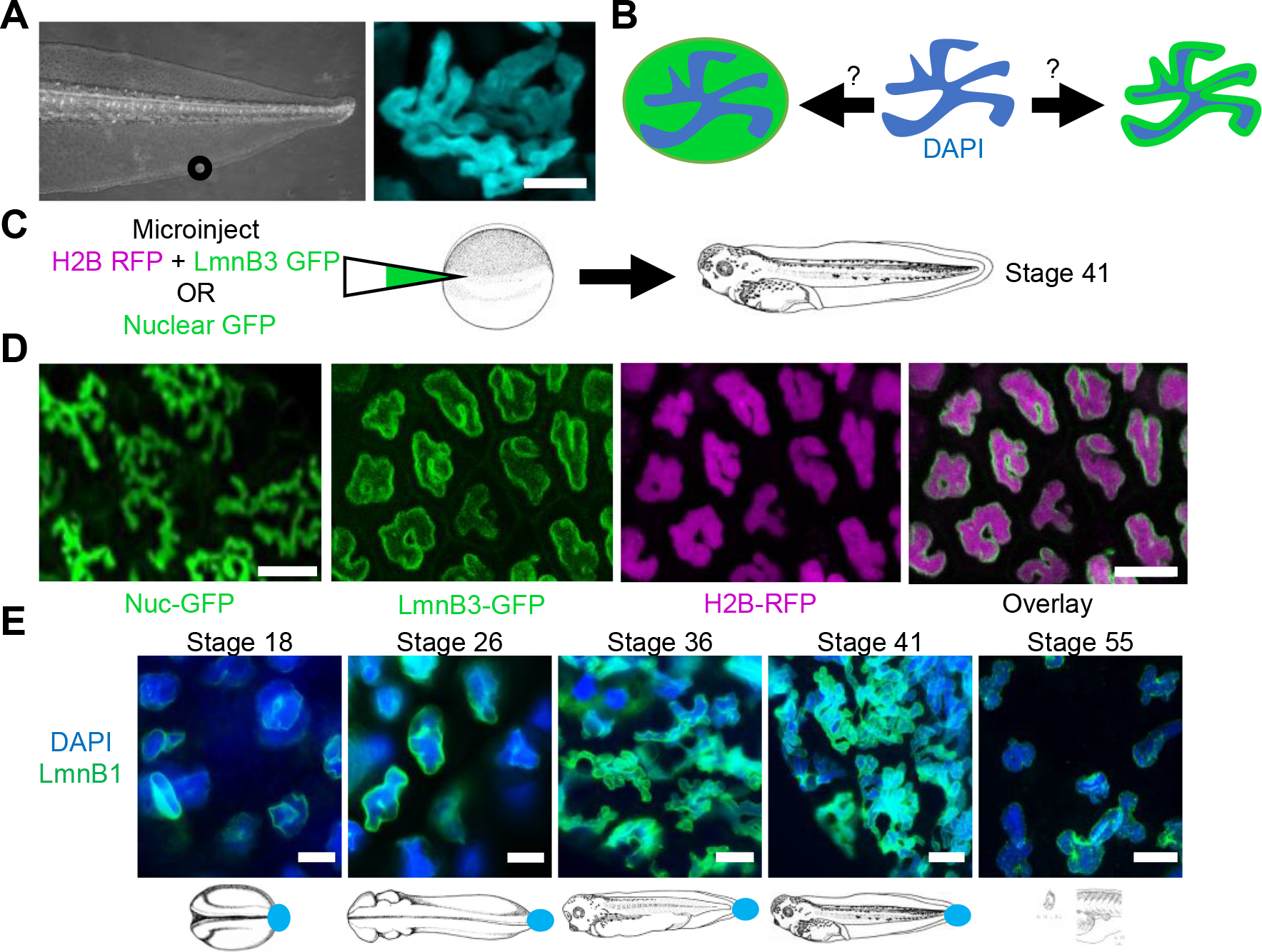
Nuclei in the tail fin of *Xenopus tropicalis* are branched. A) Bright field image of a stage 41 tadpole tail. Immunofluorescence of DAPI (cyan) in a single nucleus. The scale bar represents 5 μm. B) Two models of nuclear structure, branched chromatin in an ellipsoid nuclear compartment or branched chromatin in a branched nuclear compartment. C) Experimental design to address B). D) Fluorescent images of the nuclear lumen, periphery, and chromatin. Scale bar represents 10μm. E) Immunofluorescence of nuclear branches during development. Scale bar represents 10 μm.

We next asked what the spatiotemporal distribution of nuclear branching is during *Xenopus* development. Because injected mRNAs have a limited lifetime, we utilized immunohistochemistry to explore endogenous nuclear structure in the epidermis and other tissues through development. Tadpoles were fixed at various stages and stained for Lamin B1 to show the nuclear periphery and, DAPI to label chromatin. We find that by late neurula stages (NF stage 18), the nuclear envelope is ruffled and irregular, though the chromatin distribution is still largely ellipsoid (Fig. 1E). We note that ruffling of the nuclear envelope is found in several epidermal cell types at this stage, including secretory cells, multiciliated cells, and the goblet cells surrounding them (Fig. S1). At NF stage 22, ruffling of the nuclear envelope is more pronounced, though chromatin distribution as shown by DAPI remains ellipsoid. As the embryo enters tailbud stages, the distribution of both chromatin and the nuclear lamina becomes gradually more branched, with defined branches appearing by NF stage 26, multiple branches evident per nucleus by NF stage 35, and the most elaborate degree of branching reached by NF stage 41 (Fig. 1E). The absolute number of branches per nucleus is quiet variable, ranging dramatically from 2-13 (Fig. S1). All nuclear branching is lost shortly before the onset of tail reabsorption, and epidermal cells of the adult frog are not branched (data not shown). While non-ellipsoid nuclear structures are also visible in some other cell types, notably granulocytes/neutrophils, nuclear branching was only observed in epidermal cells. Epidermal cells of the head at stage 41 showed some minor lobulation, while nuclei of other tissues such as the heart and somites were ellipsoid (Fig. S1). The only structure in which we identified branched nuclei in outside of the tail was the surface epithelial cells covering the retina (Fig. S1).

### Branched nuclei have intact envelopes, and contain normal mitochondria, nucleoli, and marks of active enhancers

Because perturbations of nuclear morphology are associated with pathology in many cell types (Li et al., 2016; Wang et al., 2008), we asked whether epidermal cells with branched nuclei showed hallmarks of cellular damage or senescence. These could include nuclear envelope rupture, mitochondrial damage, or cell cycle exit. To assess subcellular signs of cell damage, we utilized TEM (Transmission electron microscopy) to assess nuclear envelope integrity and mitochondrial abundance. Micrographs reveal diverse nuclear structures (Fig. 2A), including clearly demarcated branched nuclei enclosed by bilayer nuclear envelopes. Upon close examination of the nuclear envelope we see that it is a continuous bilayer in cells with branched nuclei, with an average of 18 nm between the inner and outer leaflets and containing nuclear pores with a mean diameter of 60 nm (Fig. 2B). The integrity of the nuclear envelope is also supported by the even distribution of Nuclear GFP within the nuclear lumen and by the continuous distribution of LaminB1 in branched nuclei, with no evidence of leaks, partitions or ruptures (Fig. 1D, E). Cells with branched nuclei also contain numerous mitochondria with abundant cristae (Fig. 2A). These observations suggest cells with branched nuclei are not undergoing apoptotic or senescent processes that would be reflected in nuclear envelope breakdown, low mitochondrial numbers or loss of mitochondrial cristae.

**Figure 2.**
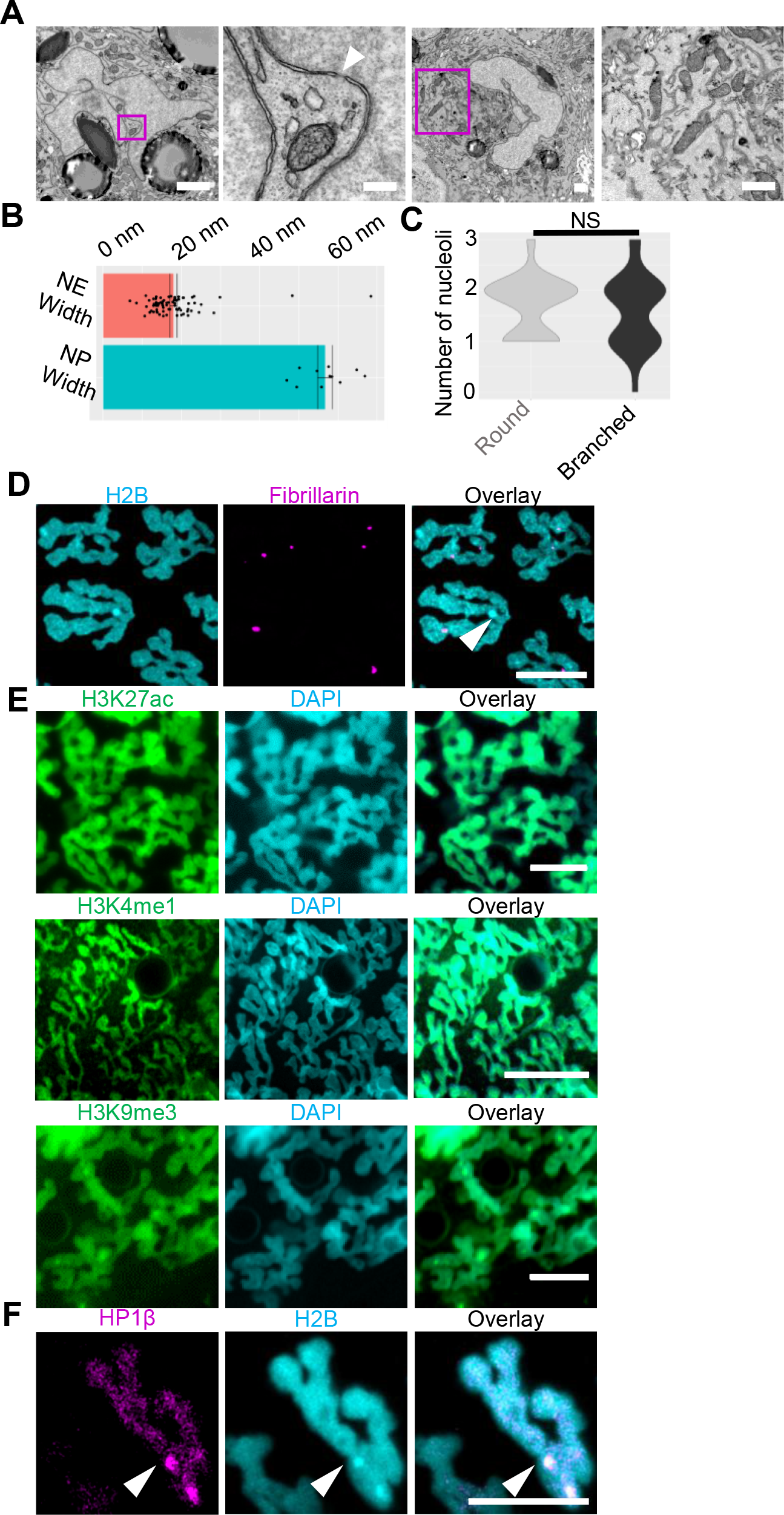
Epidermal cells with branched nuclei appear healthy and contain active enhancers. A) Transmission electron micrographs of cells with branched nuclei. Upper left panel shows a single nucleus, Scale bar represents 1μm. Magenta box indicates region depicted in second panel. Arrow head shows nuclear pore. Scale bar represents 250nm. Third panel shows a single nucleus, scale bar represents 1 μm. Magenta box indicates region depicted in fourth panel. Fourth panel shows mitochondria of a cell with a branched nucleus, with visible cristae. Scale bar represents 1 μm. B) Quantification of nuclear envelope (NE) width and nuclear pore (NP) width from TEM micrographs. C) Violin plots of the distribution of nucleoli in round (n=38 nuclei, 3 tadpoles) and branched nuclei in the tadpole (n=46 nuclei, 3 tadpoles) (p=0.49, two-tailed student’s t-test). D) Fluorescent images of H2B and nucleoli labeled by fibrillarin, scale bar represents 10 μm. E) Distribution of chromatin marks in branched nuclei. Immunofluorescence of H3K27ac (active transcription), H3K4me3 (active enhancers) H3K9me3 (heterochromatin). Scale bars represent 10 μm. F) Live image of HP1β (heterochromatin), white arrow heads indicate foci. Scale bars represent 10 μm.

TEM did reveal some atypical features in branched nuclei, including a lack of well-defined regions of perinuclear increased electron density in micrographs that would be indicative of heterochromatic regions, or of clear identifiable nucleoli (Fig. 2A). To determine if there were nucleoli present in nuclear branches we analyzed localization of the nucleolar marker fibrillarin (Brangwynne et al., 2011). We found that cells with branched nuclei did contain foci of fibrillarin (Fig. 2C), suggesting the presence of nucleoli, and that the average number of foci per nucleus did not change between branched (1.60) and unbranched nuclei (1.69) (Fig. 2D). We did note that foci of fibrillarin did not correspond to apparent foci of H2B.

To better understand whether cells with branched nuclei contain both active and inactive chromatin domains, we used immunofluorescence and live imaging to examine the distribution of histone modifications associated with active enhancers (H3K27ac and H3K4me1), and heterochromatin (H3K9me3 and HP1β). Using immunofluorescence we find H3K27ac, H3K4me1, and H3K9me3 are all distributed broadly in branched nuclei at NF stage 41 (Fig. 2E); the distribution appears uniform for H3K27ac, but H3K9me3 appeared more concentrated in foci, and H3K4me1 may be excluded from some regions of the nuclear periphery. To better characterize the distribution of heterochromatin, we used a GFP fusion of the heterochromatin binding protein HP1β (Mattout et al., 2015b). We find that HP1β-GFP broadly co-localizes with H2B (Fig. 2F). Although heterochromatin is typically enriched at the nuclear envelope, we did not observe a clear enrichment of HP1β at the nuclear periphery, however, we observe foci of HP1β-GFP fluorescence in the nucleus corresponding to foci in H2B. Taken together this suggests that active enhancers and heterochromatin are found throughout nuclear branches.

### Cells with branched nuclei have active cell cycles

In many cell types, breakdown of ellipsoid nuclear morphology is a hallmark of senescence, cell cycle dysregulation, or genomic instability (Dahl et al., 2006; Goldman et al., 2004; Schirmer et al., 2001; Wang et al., 2008) In particular, keratinocytes are known to acquire aberrant nuclear morphologies following terminal differentiation and cell cycle exit, and in premature aging syndromes (Gdula et al., 2013; McKenna et al., 2014). We therefore wanted to determine if cells with branched nuclei in the keratin-rich tadpole epidermis were undergoing an active cell cycle. We utilized immunofluorescence of Phosphorylated-Histone H3 (PH3) to mark mitotic nuclei, and Lamin B1 to mark the nuclear periphery. We find numerous examples of PH3-positive cells that retain branched nuclei (Fig. 3A). Examination of chromatin morphology in PH3-positive cells suggests that nuclei remain branched and are still enclosed by a branched nuclear envelope through prophase but form a condensed metaphase plate while the nuclear envelope breaks down. Chromatin remains condensed through anaphase. Daughter cells establish independent branching patterns and re-form the nuclear envelope at late telophase.

**Figure 3.**
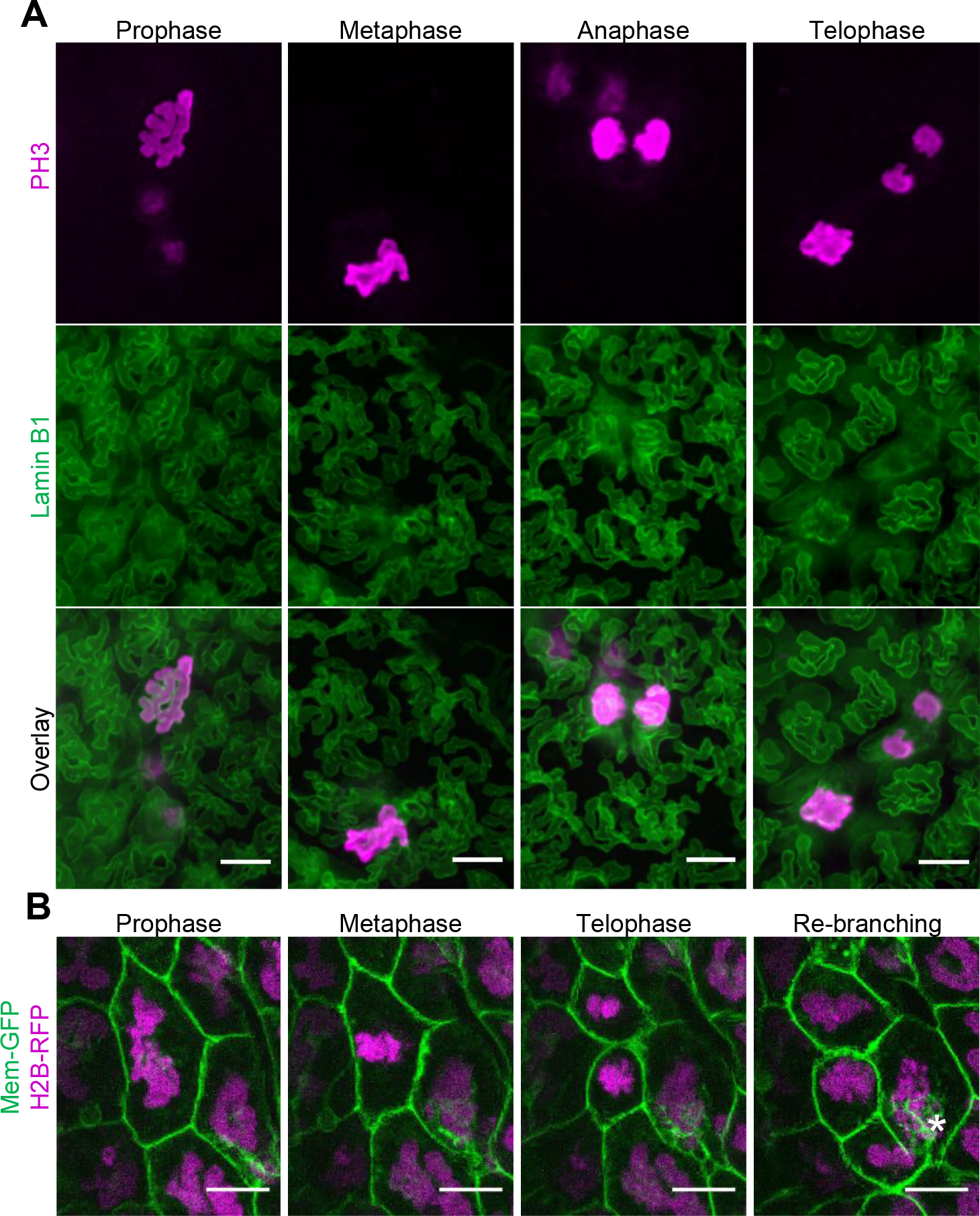
Cells with branched nuclei have active cell cycles. A) Immunofluorescence of phospho-H3 (PH3) and Lamin B1 cells in various stages of mitosis. B) Various stages of mitosis in live cells. Asterisk shows potential vesicles being from the cell. Scale bars represent 10 μm.

To better characterize nuclear envelope and chromatin dynamics through mitosis, we conducted live imaging using H2B-RFP and membrane-GFP to track individual nuclei throughout mitosis (Fig. 3B, Movie 1). These confirmed our initial observations that nuclei are initially branched, formed morphologically normal metaphase plates that segregate into two well-defined populations at anaphase, and are re-enclosed by the nuclear envelope following telophase, with the nucleus beginning to re-form branches approximately 21 minutes after cytokinesis (Movie 1). Nuclear branching patterns in daughter cells do not typically recapitulate those of the mother cell, nor do both daughters show the same branching patterns. Additionally, branches did not appear to be reabsorbed once formed after the completion of mitosis but did exhibit some dynamic motion within the branches. In nuclei not undergoing mitosis, the number and relative positions of branches can remain stable for two hours or more. Both fixed and live imaging therefore demonstrate that branched nuclei are able to undergo mitosis.

### Perturbations of Actin and not Microtubules disrupt nuclear branching

We next sought to determine what molecular mechanisms enabled nuclear branching in the fin margin. In mammalian cells, perturbations of nucleoskeleton components lead to nuclear shape deformation. These include mutations in LMNA, which lead to nuclear blebbing in progeroid syndromes (Chen et al., 2014; Dahl et al., 2006; Goldman et al., 2004; Perovanovic et al., 2016; Verstraeten et al., 2008), mutations or duplications of LMNB1 or its receptor, which disrupt nuclear flexibility and extravasation in neutrophils (Dreesen et al., 2013) perturbations of the Sun and Nesprin components of the LINC complex (Chang et al., 2015; Hatch and Hetzer, 2016; Kim and Wirtz, 2015), or alterations in the abundance, orientation, or phosphorylation of actin, which contribute to nuclear morphological disruption in HeLa cells (Ho et al., 2013; Kim and Wirtz, 2015; King and Lusk, 2016; Ramdas and Shivashankar, 2015; Webster et al., 2009; Wiggan et al., 2017; Zwerger et al., 2013). Therefore, we decided to pursue whether similar components were required for nuclear branching in the fin margin.

First, we observed actin localization in cells with branched nuclei. We found no apparent bias of actin localization to tips or bases of branches (Fig. 4A). To determine if actin filaments were necessary for nuclear branches we incubated stage 41 tadpoles with Latrunculin B (Lat B), which disrupts actin filament formation which Lat B has been found to disrupt other actin-dependent processes in *Xenopus* at these non-lethal doses (Lee and Harland, 2007). To monitor the effect of this inhibitor on actin filaments and nuclear morphology, we injected embryos at cleavage stages with mRNAs encoding H2B-RFP and the actin binding protein Utrophin-GFP. We find that treatment with Lat B results in breakdown of the actin cytoskeleton beginning at 25 minutes post treatment. At this time, foci of Utrophin-GFP were visible (Movie 2, Fig. 4B). Nuclear branches were gradually lost after actin destabilization and were lost more slowly in nuclei that initially had more numerous or complex branches. Nuclear branches reformed after wash-out of Lat B. Actin filaments visualized by LifeAct began to be visible 25 minutes after Lat B removal along with some nuclear deformation, similar kinetics to what was observed for the loss of actin filaments. By 125 minutes after Lat B removal new branches were fully formed although the branching patterns were not conserved relative to their initial pre-treatment distribution (Fig. 4C).

**Figure 4.**
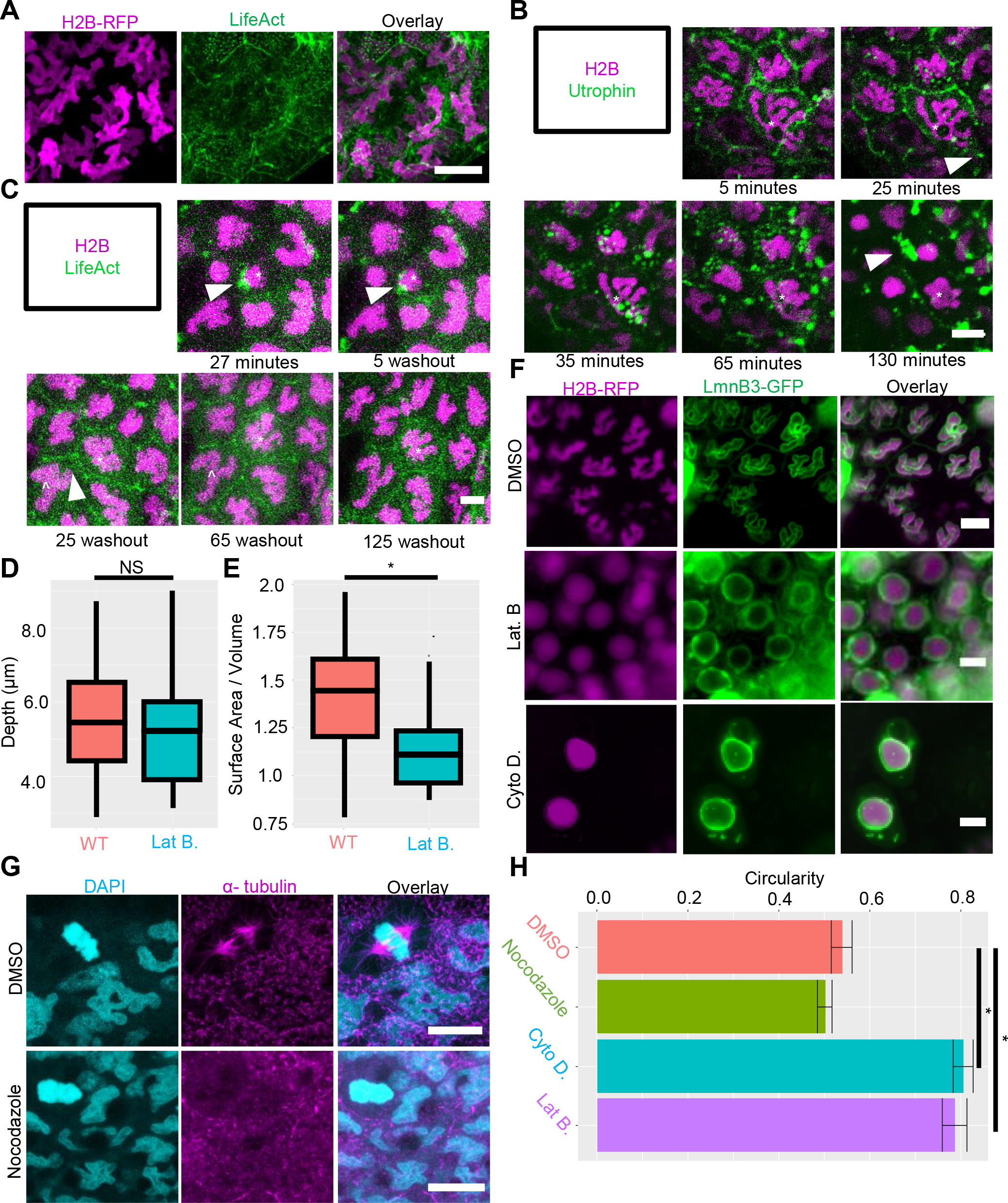
Perturbations of Actin but not microtubules disrupt nuclear branching. A) Actin localization in cells with branched nuclei, H2B (magenta) and LifeAct (green) Scale bar = 10 μm. B) Latrunculin B treatment causes loss of actin filaments (utrophin-GFP, green), and nuclear branches (H2B, magenta). White arrow heads show depolymerized actin, asterisk denotes a single nucleus. Times denote length of treatment. Scale bar = 10 μm. C) Nuclear branches (H2B, magenta) and actin filaments (LifeAct, green) reform after Lat B wash-out. White arrow heads show changes in actin. Asterisk and carrot show single nuclei. Scale bar = 10 μm. D) Nuclear depth (μm) of Wildtype (n=14 nuclei, 3 tadpoles) and Lat B treated tadpoles (n=17 nuclei, 4 tadpoles) (E) Nuclear surface area / volume ratios in Wildtype (n=21 nuclei, 3 tadpoles) and Lat B treated tadpoles (n=19 nuclei, 3 tadpoles) (p<0.05, one-tailed student’s t-test) F) Treatment with Cytochalasin D (Cyto D) and latrunculin B (Lat B) disrupt nuclear branches, branches remain intact in DMSO vehicle control. Scale bars =10 μm. G) Nocodazole treatment disrupts microtubules, but not nuclear branches. Scale bars =10 μm. H) Quantification of nuclear circularity in actin and microtubule drug treatment, Cyto D (n=50; 7 tadpoles) and Lat B (n=45; 6 tadpoles) significantly increase circularity compared to DMSO (n=168; 10 tadpoles), Nocodazole (n=90; 3 tadpoles) had no change compared to DMSO (p<0.01, one-way ANOVA and Tukey’s post-hoc, error bars are s.e.m.) C) Mosaic CRISPR/Cas9 knock-out of lamin B1, and dominant negative lamin B1 disrupt nuclear branches. Scale bars represent 10 μm. D) Quantification of nuclear circularity in lamin B1 perturbed nuclei. Whole animal (n=86 nuclei; 7 tadpoles) and epidermal only knockouts (n=51 nuclei; 6 tadpoles), and laminB1 dominant negative (n=50; 5 tadpoles) increase circularity significantly relative to wildtype (n=82 nuclei; 7 tadpoles) (p<0.01, ANOVA and Tukey’s post hoc, error bars are s.e.m.). Scrambled CRISPR/Cas9 guide (n=40; 5 tadpoles) does not change nuclear circularity relative to wildtype.

Because actin has known roles in compressing nuclei (Versaevel et al., 2012; Vishavkarma et al., 2014; Wiggan et al., 2017) and nuclear branches are in an extremely flattened epithelium, we next measured changes in nuclear depth and surface area -to- volume ratios of nuclei with intact and Lat B perturbed actin networks. Both WT (wild type) and Lat B-treated tadpoles had comparable nuclear depths (5.6 and 4.1 μm respectively) (Fig. 4D). However, the nuclear surface area -to- volume ratio decreased from 1.405 in WT to 1.134 in Lat B treated animals (Fig. 4E). Nuclear volume also decreased by approximately 7.5% and the surface decreased by approximately 25.4% in Lat B treated animals (Data not shown). Together this suggests that loss of branches decreases the amount of nuclear membrane (surface area) relative to the volume. However, the lack of change in nuclear depth suggests that actin is not suppling a compressive force causing branches to form, but rather pushing, or pulling forces.

To confirm that loss of nuclear branches was not specific to Lat B treatment we utilized Cytochalasin D (Cyto D), which inhibits actin polymerization and has also been shown to disrupt other actin-dependent processes in *Xenopus* (Lee and Harland, 2007). Treatment with either Cyto D or Lat B results in a rapid loss of nuclear branching in the fin margin, as revealed by LamnB3-GFP and H2B-RFP (Fig. 4F).

We next asked whether microtubules contributed to nuclear branches, as they have been shown to play a role in maintaining nuclear morphology(Tariq et al., 2017). We utilized the microtubule polymerization inhibitor nocodazole at non-lethal doses (Dutta and Kumar Sinha, 2015). While nocodazole treatment noticeable disrupts spindle formation in tadpoles, it does not affect nuclear morphology relative to DMSO-treated controls (Fig. 4G).

We quantified changes in nuclear morphology for all cytoskeleton perturbations using the circularity measurement on ImageJ (see methods, Fig. S2). We found there was a statistically significant increase in epidermal nuclear circularity when tadpoles were treated with either Cyto D or Lat B, but not Nocodazole (Fig. 4H). These results indicate that nuclear branches require intact actin filaments for their maintenance, but not polymerized microtubules.

### LaminB1 is necessary for nuclear branches

We next asked whether nuclear branching relies on specific components of the nuclear lamina. Modulation of nuclear lamina components has been shown to regulate tissue elasticity in mammals: greater amounts of Lamin A contribute to stiffer tissue, while Lamin B is critical for nuclear envelope flexibility in neutrophils (Mattout et al., 2015a; Mattout et al., 2015b; Peric-Hupkes et al., 2010; Perovanovic et al., 2016; Solovei et al., 2013; Towbin et al., 2012). *Xenopus tropicalis* contain one Lamin A/C homolog, as well as three Lamin B homologs: Lamin B1, Lamin B2 and the germline-specific Lamin B3 (Session et al., 2016). We first sought to determine whether any of these components were preferentially enriched or depleted in fin marginal cells containing branched nuclei. To this end, we isolated fin margin tissue or whole embryo tissue, and quantified expression of *lmnb1*, *lmnb2*, and *lmna* using qRT-PCR (Fig. S2). We find that expression of *lmnb1* is significantly upregulated in the fin margin relative to the whole embryo (2.6-fold increase), whereas *lmnb2* and *lmna* are unchanged. To determine whether this upregulation reflected a functional role for *lmnb1* in the fin margin or in nuclear branching, we used CRISPR/Cas9 to create mutations in *lmnb1* by co-injecting gene-specific sgRNAs together with humanized Cas9 protein in F0 tadpoles (Bhattacharya et al., 2015; Nakayama et al., 2013). To track nuclear and cell morphology, we again co-injected these embryos with mRNAs encoding H2B-RFP and Membrane-GFP. We used high-resolution melt analysis to confirm gene-specific mutations (Fig. S2). Upon analyzing nuclear morphology in F0 tadpoles at stage 41, we find that only *lmnB1* mutant embryos (*Lmnb1* CRISPR) have markedly reduced branching, instead exhibiting crescent or elongated obloid shapes (Fig. 5A). This effect is confined to *lmnB1* mutants and is not induced by injection of a scrambled version of the *lmnB1* sgRNA (Scrmbl) (Fig. 5A).

**Figure 5.**
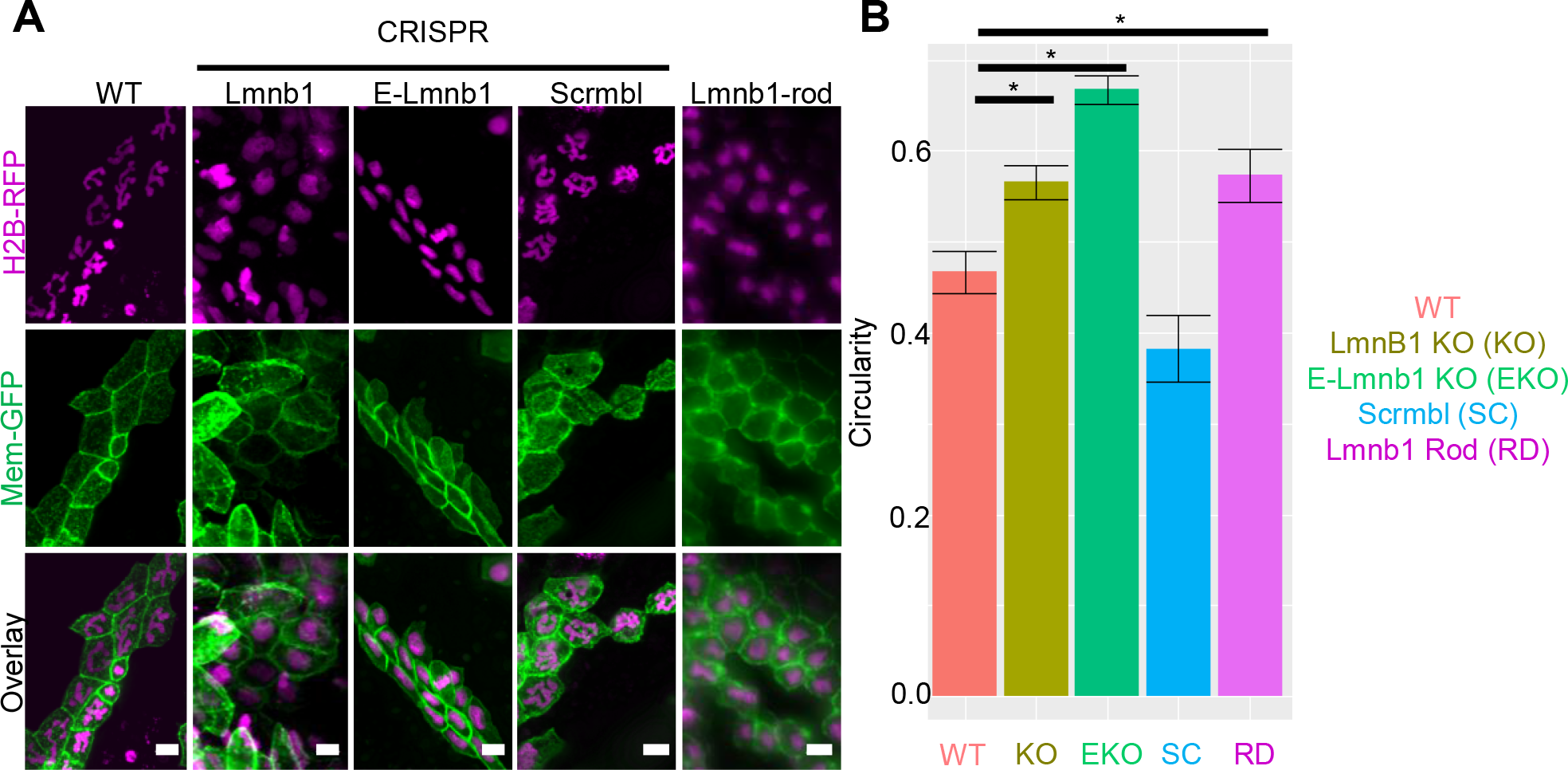
LaminB1 is necessary for nuclear branches. A) Mosaic Crispr Cas9 knock-out of lamin B1, and dominant negative lamin B1 disrupt nuclear branches. Scale bars represent 10 μm. B) Quantification of nuclear circularity in lamin B1 perturbed nuclei. Whole animal (n=86 nuclei; 7 tadpoles) and epidermal only knockouts (n=51 nuclei; 6 tadpoles), and laminB1 dominant negative (n=50 nuclei; 5 tadpoles) increase circularity significantly relative to wildtype (n=82 nuclei; 7 tadpoles) (p<0.01, ANOVA and Tukey’s post hoc, error bars are s.e.m.). Scrambled CRISPR/Cas9 guide (n=40 nuclei; 5 tadpoles) does not change nuclear circularity relative to wildtype.

To confirm that these effects were intrinsic to epidermal cells, and not secondary to any effect of whole-embryo perturbation, we next targeted our injections specifically to the ventral animal blastomeres at the 8-cell stage, which give rise to the epidermal lineage (Bauer et al., 1994; Moody, 1987). These epidermal-only *lmnB1* mutant embryos (E- Lmnb1 CRISPR) also exhibited reduced nuclear branching in the fin epidermis (Fig. 5A). As a further confirmation, we generated a *Xenopus* form of dominant-negative Lamin B1, following the domain structure used in mammals (Goldman et al., 2004). This dominant negative LaminB1 (Lmnb1-rod) contains only the rod domain, which is thought to disrupt the lamin network and LINC complex interactions when overexpressed (Goldman et al., 2004). We co-injected this into epidermal blastomeres at the 8-cell stage, together with H2B-RFP and Membrane-GFP. Epidermal cells injected with Lmnb1-rod also exhibited reduced nuclear branching at stage 41 (Fig. 5A). We quantified nuclear circularity in LmnB1 perturbed tadpoles and found that epidermal cells in LmnB1 CRISPR, E-LmnB1 CRISPR, and Lmnb1-rod tadpoles had statistically significant increase in nuclear circularity, where there was no difference between WT and Scrmbl tadpoles (Fig. 5B). Together these results argue that nuclear branching depends on functional Lamin B1 in the epidermis.

### Nuclear morphology arises independently from swimming motions and contributes to fin morphology

One potential source of nuclear morphological variation derives from the physical forces exerted against the nucleus either by intracellular compression, such as through perinuclear actin, or by extracellular compression, such as that exerted by endothelial cells during neutrophil extravasation. The tadpole fin encounters a unique extracellular force profile as it undergoes swimming movements. We therefore tested if the mechanical forces sustained by the fin from swimming were required for nuclear branching in the fin. To this end, we utilized dorsal posterior explants at the neurula stage. These explants give rise to tails with fins that lack muscle (Tucker and Slack, 2004). We then compared the circularity of nuclei in the fin margin of stage matched tadpoles and nuclei in the fin of explants and found no change in circularity (Fig. 6A). This suggests that the mechanical forces from swimming are not necessary to induce nuclear branching.

**Figure 6.**
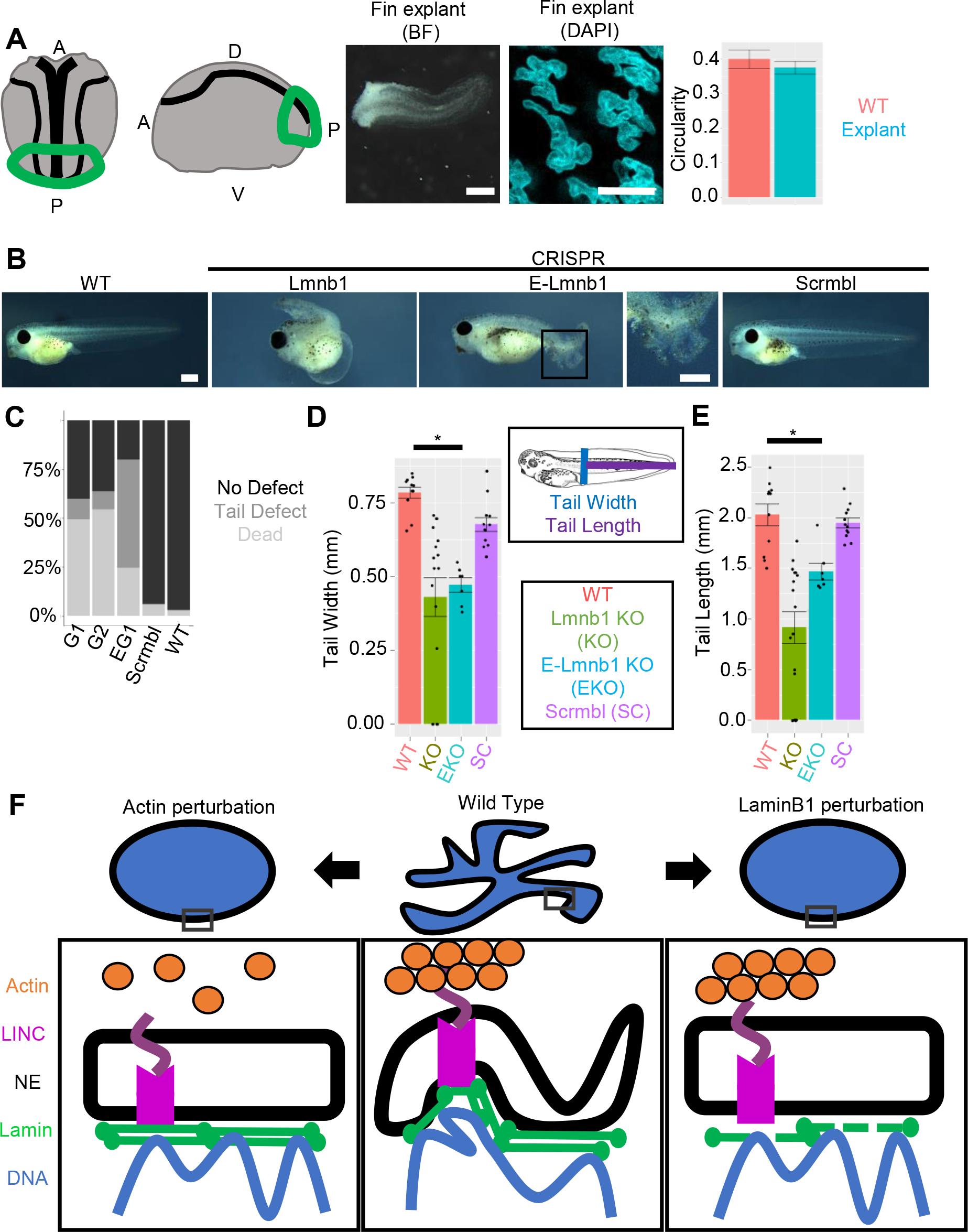
Nuclear morphology in arises independently from swimming motions and contributes to fin morphology. A) Dorsal posterior explants develop a stationary tail (scale bar indicates 1 mm) which retains nuclear branching (Scale bars represent 10 μm). Circularity is unchanged between explants (n= 86 nuclei; 5 explants) and stage matched tadpoles (n= 32 nuclei; 4 tadpoles) (p>0.05, ANOVA and Tukey’s post-hoc, error bars are s.e.m.). B) Stage 41 tadpoles with and without Lmnb1. Boxed area of E-Lmnb1 tadpole shown. Scale bars indicate 1mm. C) Tadpole phenotypes G1 = Whole embryo *lmnb1* CRISPR with guide 1, G2 = Whole embryo *lmnb1* CRISPR with guide 2, 8G1 = Epidermal *lmnb1* CRISPR with guide 1 (G1 n= 409 tadpoles; 5 clutches, G2 n=22 tadpoles; 1 clutch, 8G1 n= 155 tadpoles; 3 clutches, scrmbl n= 149 tadpoles; 3 clutches, WT n= 201 tadpoles; 4 clutches). D) Tail width of tadpoles are significantly decreased in *lmnb1* perturbed tadpoles E) Tail length of tadpoles are significantly decreased in *lmnb1* perturbated tadpoles (For D-E; WT n=11, Lmn B1 KO n= 16, E Lmn B1 KO n=6, Scrmbl n=12, p<0.01 one-way ANOVA and Tukey’s post hoc). F) Perturbations of actin filaments and LaminB1 alter nuclear morphology.

We concluded by investigating the relationship between nuclear branching and development. We observed that tadpoles with *lmnb1* mutations had tail defects, sloughing of the fin epidermis, were inefficient swimmers, and developed edema, likely due to decreased locomotion (Fig. 6B, C). We found that nuclear morphology and fin morphology were correlated; in *imnb1* sgRNA-injected tadpoles that had no tail defect, we did not observe changes in nuclear morphology (Fig. S2). Tails were statically significantly shorter and narrower in tadpoles with *lmnb1* mutations compared to wild type (Fig. 6D, E). This suggests that *lmnb1*-dependent nuclear branching may be necessary for proper tail formation and function but does not rule out the possibility that the contribution of the function of LaminB1 on gene regulation affects tail formation.

## Discussion

### *Xenopus* epidermal branching as a model for extreme variation in healthy nuclear morphology

Across tissues and species, nuclear morphology is generally ellipsoid. There are very few cases of healthy epithelial cell types with highly irregular nuclear morphologies; the mandibular gland epithelium of the wax moth *Ephestia kuehniella* is a notable example, which is exhibits a branched nuclear morphology similar to the morphology of the tadpole fin (Buntrock et al., 2012). Here we describe an epithelial epidermal tissue, the *Xenopus* tadpole fin margin, which exhibits a highly branched nuclear morphology. In this epithelial tissue, a heterogenous population of cell types including secretory cells, goblet cells and multiciliated cells displays highly irregular branched nuclear morphology. The degree of nuclear branching we describe is more extreme than in most other instances of nuclear morphological changes among both healthy and diseased vertebrate cells. Like neutrophils, these fin margin cells develop branched nuclear morphology over the course of development, but unlike neutrophils appear to decrease branching to some degree as tadpoles age.

*Xenopus* has long served as a model organism for nuclear morphology, including molecular and cell biological characterization of nuclear envelope components and the cell biological consequences of their perturbation. Overexpression of specific Lamin components has been observed to alter both nuclear shape and size in oocyte nuclei from *Xenopus laevis*, and nuclear size scaling in early *Xenopus* embryos is dependent on cytoplasmic volume as well as the nuclear transport factors Importin a and NTF2 (Good et al., 2013; Jevtić and Levy, 2015; Jevtić et al., 2015; Levy and Heald, 2010). More recently, *Xenopus* has served as a source model for proteomic studies of nuclear composition (Wühr et al., 2015). Morphological variation in nuclei later in embryogenesis has not been examined in depth. We find that nuclear branching begins late in neurulation, well after the initial specification of epidermal fate but similar to the stage when multiciliated cells begin to undergo apical emergence (Sedzinski et al., 2016). Nuclear branching is dramatically elaborated as the tail elongates, and by late tailbud stages highly branched nuclei are found both in multiciliated cells and their goblet cell neighbors. Our data suggest that nuclear branching is a general property of the fin margin epithelium in *Xenopus*. The extremity of morphological variation observed suggests that these nuclei may represent a valuable model for nuclear diversity: they are easily imaged, in a whole organism system that is easily modulated both genetically and through small molecules.

### Branched nuclei do not interfere with cell health or mitosis

*Xenopus* epidermal fin cells exhibit branched nuclear morphologies while maintaining an active cell cycle. This is in contrast to many cases of non-ellipsoid nuclear morphologies, which occur in post mitotic-cells. In the epidermis, keratinocytes are known to undergo nuclear flattening and to acquire irregularities in their nuclear envelope after cell cycle exit, and these become more extreme with aging or in specific disease scenarios (Yang et al., 2011). Among actively-cycling cells, nuclear morphological perturbations such as blebbing or nuclear ruffling are common in cancer cells but very infrequent in healthy cell types (Denais and Lammerding, 2014; Fu et al., 2012; Pillay et al., 2013; Shah et al., 2013). We find that nuclear branching is common in mitotic epidermal cells in the *Xenopus* tail fin, with rapid collapse of nuclear branching approximately 7 minutes before metaphase and re-formation of nuclear branches becoming apparent 21 minutes after cytokinesis (Movie 1). Branched nuclei do appear to undergo complete nuclear envelope breakdown, and we have not found evidence of karyomeres or chromosome-specific nuclear envelopes as are seen in the early mitoses of zebrafish and *Xenopus* (Lemaitre et al., 1998; Schoft et al., 2003). Following mitosis, the branched structure formed by the two daughter cells are distinct and do not faithfully recapitulate the mother cell’s nuclear morphology.

In cells with branched nuclei we find that both active enhancers and heterochromatin are distributed continuously throughout the nucleus. Normally, regions of chromatin that are transcriptionally repressed are associated with the nuclear lamina and the periphery of the nucleolus (Mattout et al., 2015a; Mattout et al., 2015b; Peric-Hupkes et al., 2010; Perovanovic et al., 2016; Solovei et al., 2013; Towbin et al., 2012). In TEM of branched epidermal nuclei we see no increase in electron density around the nuclear periphery, nor do we find evidence of increased electron density representing a nucleolus in most nuclei. However, we found that cells with branched nuclei did contain foci of fibrillarin, a component of the nucleolus, in regions distinct from the densest chromatin as represented by H2B fluorescence. Interestingly we do see puncta of increased intensity of HP1β and H3K9Me3 but have not yet seen corresponding regions of increased density on TEM. Taken together this suggests that cells with branched nuclei may partition heterochromatin without anchoring these regions to the nuclear lamina. Future research will examine the organization of heterochromatin in branched nuclei, and the organization of specific chromatin domains within these nuclei.

### Nuclear branching in tail fin is dependent on nucleoskeleton components

Previous work has shown a role for the nuclear lamina, LINC complex, and cytoplasmic actin in the shaping of the nucleus (Chen et al., 2014; Hatch and Hetzer, 2016; Hatch et al., 2013; Ho et al., 2013; Kim and Wirtz, 2015; King and Lusk, 2016; Lammerding et al., 2006; Ramdas and Shivashankar, 2015; Webster et al., 2009; Wiggan et al., 2017). Here we have shown that both an intact actin network and Lamin B1 are necessary to maintain nuclear branches. Our working model therefore suggests that actin and LaminB1 filaments serve to maintain nuclear branches by stabilizing curvature across the nuclear envelope. Loss of either filamentous actin or LaminB1 results in a broken bridge across the nuclear envelope disrupting local curvature without envelope blebbing (Fig. 6F). Additional experiments will be needed to clarify how the loss of Lamin B1 may indirectly affect the localization of other Lamin sub-types or binding partners affect nuclear branching.

Actin is known to play a role in compressing the nucleus with stress fibers during migration or passage through narrow openings (Versaevel et al., 2012; Vishavkarma et al., 2014; Wiggan et al., 2017). In laminopathies nuclei lose rounded morphologies and adopt more irregular architectures. Previous studies have also shown that gaps in the nuclear lamina allow blebbing (Hatch et al., 2013). Conversely, the fin margin nuclei appear to have a fully functional lamina network with no apparent gaps. There was no obvious localization of actin to the base or tips of nuclear branches indicative of actin pushing or pulling on the nucleus, but, loss of actin caused nuclei to increase in circularity, suggesting that by some other mechanism they contribute to nuclear branching. We also found that the loss of actin did not increase nuclear depth suggesting that the extra-cellular matrix or other force transmitting molecules cause the flattening of this tissue. We did find that there was a decrease in nuclear surface are to volume ratio when f-actin was lost, as well as a modest decrease in total nuclear volume. These observations both suggest that nuclear envelope distribution, and possibly quantity, are closely linked to f-actin in these nuclei. While it is clear f-actin is necessary to maintain nuclear branches, it is unclear how nuclear actin or actin binding proteins contribute to maintenance, and establishment of nuclear branches. Previous studies have shown a relationship between cell-spreading and nuclear actin polymerization raising the possibility that in this flattened epithelium nuclear actin may contribute to nuclear branch formation (Keeling et al., 2017; Plessner et al., 2015). Another possibility that we have not yet been able to test explicitly is the role for intra-nuclear actin filaments (Baarlink et al., 2017; Kalendová et al., 2014; Oda et al., 2017), which may also contribute to nuclear branching. Out time-lapse movies show that nuclear morphological change tracks closely in time with cytoplasmic f-actin disruption. We therefore favor the hypothesis that cytoplasmic f-actin is critical to nuclear morphology, but intranuclear actin may also contribute to the formation or stabilization of branches.

### A potential biological function for nuclear branching

Perturbations of nuclear branching have deleterious effects on the formation of the fin and consequently on its downstream function. While we have been able to show that specific nucleoskeletal components are required for nuclear branching, the ultimate role of branched nuclear morphologies in tail fin cell and tissue function remains open. Nuclear branching may play a role in genomic organization or gene regulation, as discussed above, or in fin biomechanics.

The thin epithelium of the tadpole fin is made up of flattened epidermal cells that overlie a mesenchymal core. Its specialized cell biological and biophysical properties allow rapid regeneration and sinusoidal swimming movements (Tucker and Slack, 2004). To accommodate this structure, a flattened nuclear structure would be advantageous, and nuclear branching could impart biophysical properties necessary for tissue function. The elastic modulus of the nucleus has been shown to be different than that of the cytoskeleton. The irregular nuclear structure could aid in creating a more uniform elastic modulus of the tissue, as opposed to localized regions of differential stiffness (Guilak et al., 2000; Kha et al., 2004; Pajerowski et al., 2007). The requirement of Lamin B1 to maintain nuclear branches suggests that nuclear branching could be modulating tissue stiffness (Kha et al., 2004; King and Lusk, 2016; Pajerowski et al., 2007; Swift et al., 2013; Verstraeten et al., 2008; Zwerger et al., 2013).

In conclusion, we have shown that the fin epithelium of the *Xenopus tropicalis* tadpole tail contains a heterogenous population of cells that have branched nuclear structures. These cells with branched nuclei are healthy and have active cell cycles. Additionally, we have shown that nuclear branching depends on an intact actin network and Lamin B1. We determined that forces incurred from swimming are not necessary to induce nuclear branches, however, loss of nuclear branching through *lmnb1* mutations decreases swimming efficiency and impede tail and fin development. These cells offer a novel system to study extreme nuclear morphological variation in a healthy tissue.

## Materials and Methods

### Ovulation, in vitro fertilization, and rearing of embryos

Use of *Xenopus tropicalis* was carried out under the approval and oversight of the IACUC committee at UW, an AALAC-accredited institution. Ovulation of adult *X. tropicalis* and generation of embryos by in vitro fertilization according to published methods (Khokha et al., 2002; Sive et al., 2010). Fertilized eggs were de-jellied in 3% cysteine in 1/9x modified frog ringer’s solution (MR) for 10-15 minutes. Embryos were reared as described (Khokha et al., 2002). Staging was assessed by Nieuwkoop and Faber (Nieuwkoop and Faber, 1994).

### mRNA synthesis and injections

DNA plasmids were linearized at appropriate restriction sites (Table 1) and mRNA was transcribed with Sp6 mMessage mMachine kits (Ambion). mRNAs were injected into embryos at the 1-8 cell stage, depending on experiment, with doses indicated in Table 1.

**Table 1.**
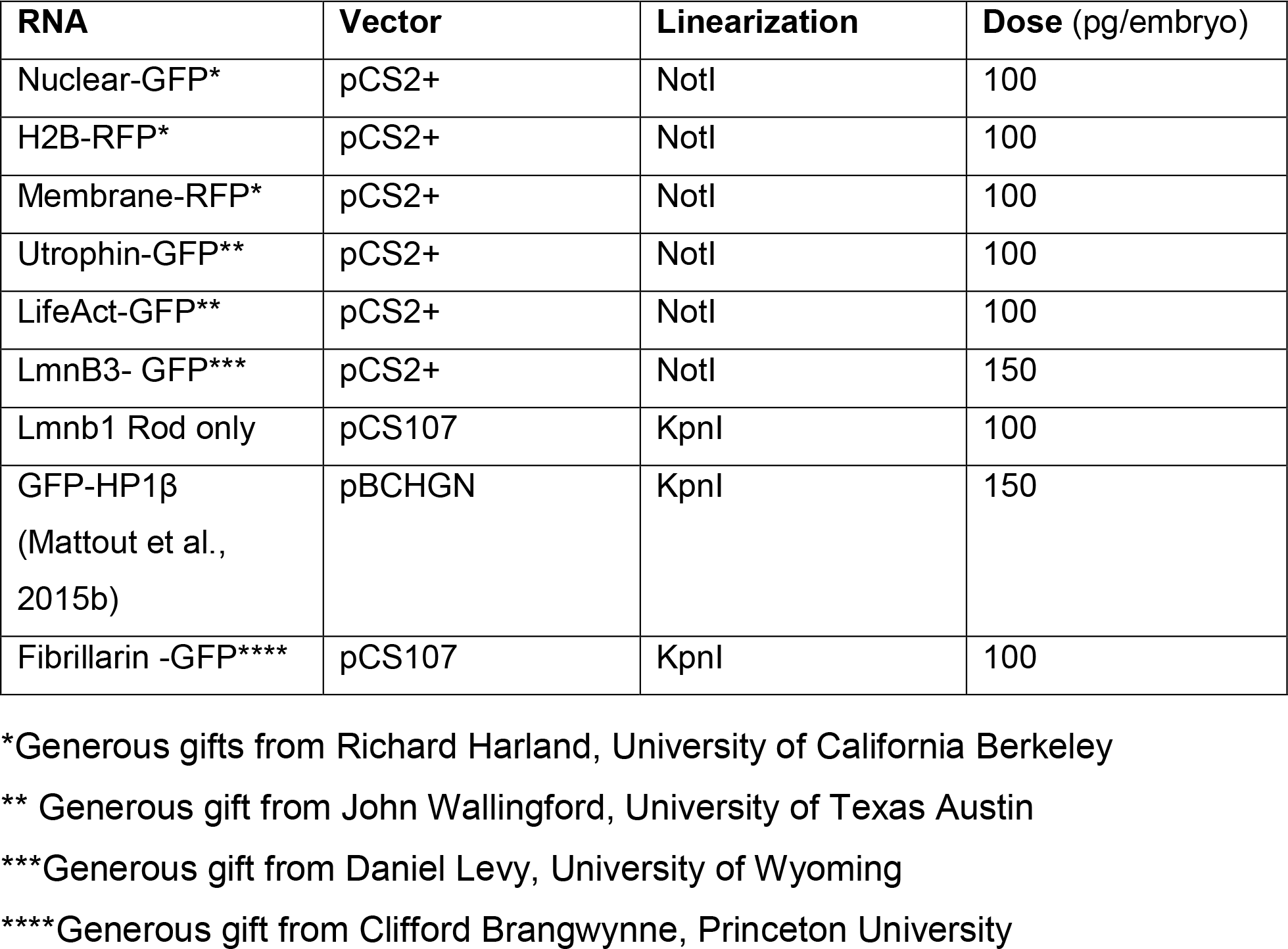

| RNA | Vector | Linearization | Dose (pg/embryo) |
| --- | --- | --- | --- |
| Nuclear-GFP* | pCS2+ | NotI | 100 |
| H2B-RFP* | pCS2+ | NotI | 100 |
| Membrane-RFP* | pCS2+ | NotI | 100 |
| Utrophin-GFP** | pCS2+ | NotI | 100 |
| LifeAct-GFP** | pCS2+ | NotI | 100 |
| LmnB3- GFP*** | pCS2+ | NotI | 150 |
| Lmnb1 Rod only | pCS107 | KpnI | 100 |
| GFP-HP1β<br>(Mattout et al.,<br>2015b) | pBCHGN | KpnI | 150 |
| Fibrillarin -GFP**** | pCS107 | KpnI | 100 |

\*Generous gifts from Richard Harland, University of California Berkeley
\*\* Generous gift from John Wallingford, University of Texas Austin
\*\*\*Generous gift from Daniel Levy, University of Wyoming
\*\*\*\*Generous gift from Clifford Brangwynne, Princeton University

### Immunohistochemistry

*X. tropicalis* embryos were fixed for 20 minutes in MEMFA at room temperature. Embryos were permeabilized by washing 3X 20 minutes in PBS + 0.01% Triton x-100 (PBT). Embryos were blocked for 1 hour at room temperature in 10% CAS-block (Invitrogen #00-8120) in PBT. Then embryos were incubated in primary antibodies (see table below) in 100% CAS-block overnight at 4°C. Embryos were then washed 3X 10 minutes at room temperature in PBT and re-blocked for 30 minutes in 10% CAS-block in PBT. Secondary antibodies (see table below) were diluted in 100% CAS-block and incubated for 2 hours. Embryos were then washed 3X 20 minutes in PBT. Whole embryos or isolated tails were mounted on slides in Vectashield containing DAPI (Vector Laboratories #H-1500). Images were acquired with a Lecia DM 5500 B and ORCA-flash 4.0LT camera.

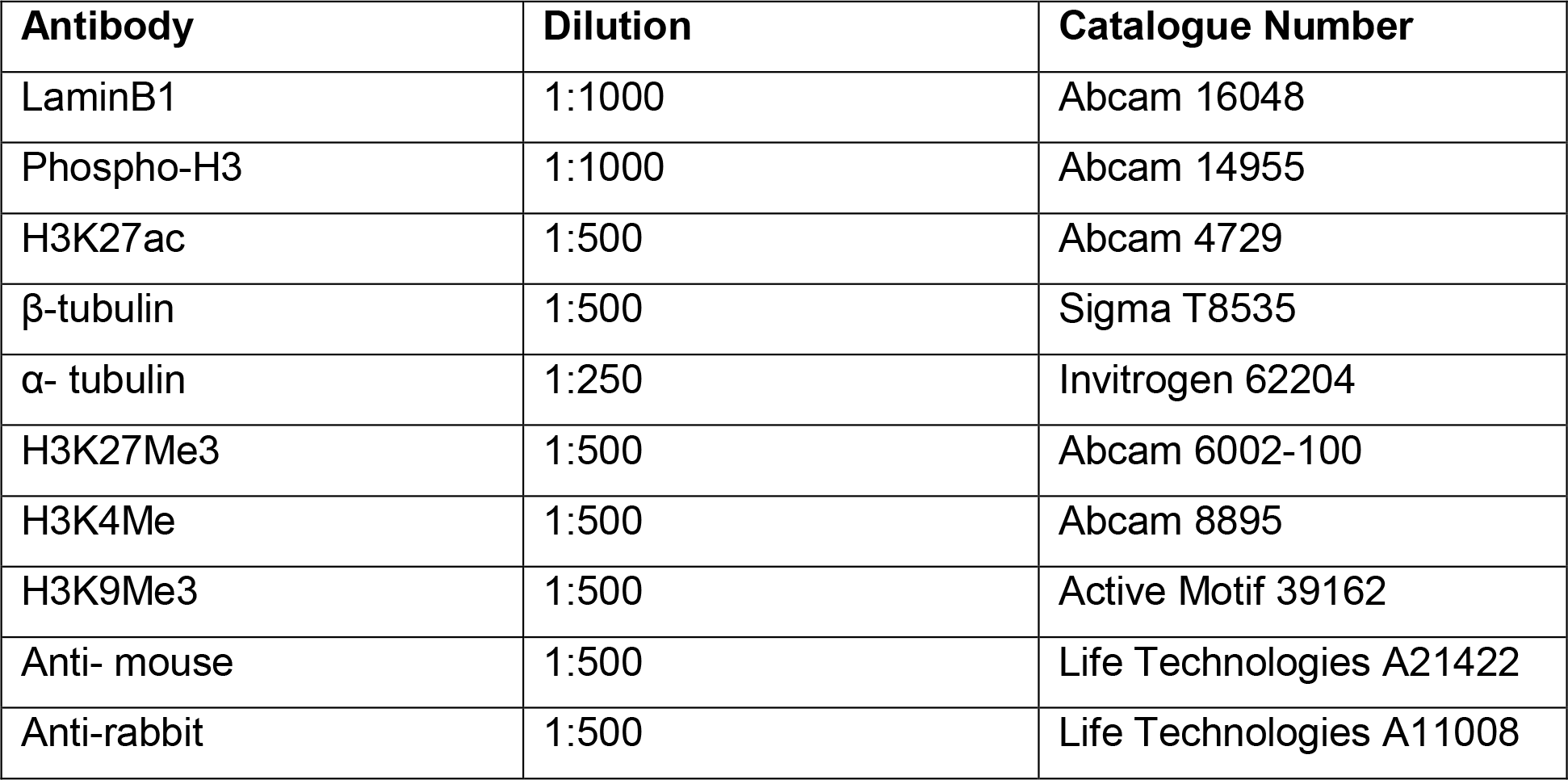

### Quantification of the number of nuclear branches

Branches were counted as the number of termini of the nucleus (Fig. S1), Branches were counted from images of tadpoles with nuclear markers of H2B, DAPI, or Nuclear localized GFP.

### Live imaging conditions

Tadpoles were imaged sedated in 0.01% tricaine in 1/9^th^ MR. Tadpoles were mounted for imaging as previously described (Kieserman et al., 2010; Wallingford, 2010) with the following modifications for Actin and Lamin B1 perturbation (Fig. 4,5): A perimeter of vacuum grease was made on a glass slide. A tadpole was placed in the center of the vacuum grease perimeter with several drops of media containing drug. A glass cover slip was gently pressed into the vacuum grease perimeter over the tadpole. Images were acquired with a Lecia DM 5500 B. Mitosis and Actin perturbation movies were acquired with a Zeiss 880. Gross tadpole morphologies were acquired with a Lecia M205 FA.

### Transmission electron microscopy

Stage 41 tadpoles were fixed in 2.5% glutaraldehyde/0.1M sodium cacodylate buffer. Samples were washed 4 times in sodium cacodylate buffer, postfixed in osmium ferrocyanide (2% osmium tetroxide/3% potassium ferrocyanide in buffer) for 1 h on ice, washed, incubated in 1% thiocarbohydrazide for 20 min, and washed again. Samples were washed and *en bloc* stained with 1% aqueous uranyl acetate overnight at 4°C. Samples were finally washed and *en bloc* stained with Walton’s lead aspartate for 30 min at 60°C, dehydrated in a graded ethanol series, and embedded in Durcupan resin. Serial sections were cut at 60 nm thickness and viewed on a JEOL-1230 microscope with an AMT XR80 camera (Giarmarco et al., 2017).

### qPCR

Total RNA was isolated from embryos (3-5 per experiment) or fin margin (15-20 per experiment) (Sive et al., 2010). RNA was treated with DNAse 1 (Invitrogen #18068015). cDNA was synthesized using SuperScript III first strand synthesis kit (Intivrogen #18080-051). Quantitative PCR analysis was preformed using BioRad iCycler PCR machine, iQ Sybr Green mix (BioRad #1708862) and analysis software.

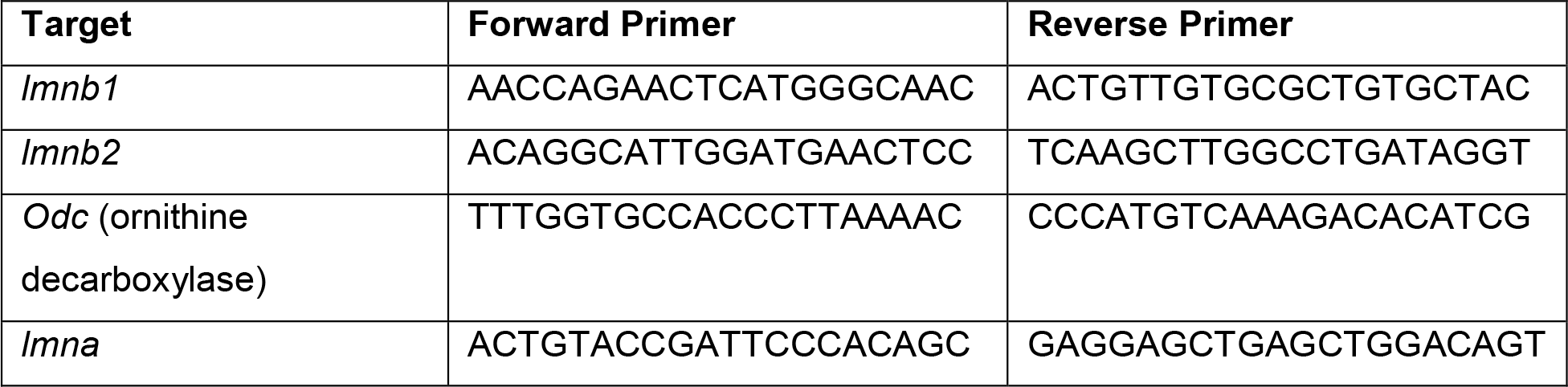

### Nuclear circularity quantification

A gaussian blur was applied to all images in a data set and a threshold was applied to images. Particles were selected using FIJI (ImageJ) and manually refined. Particles were discarded if the whole nucleus was not in the field of view, if a partial particle was selected based on the original image, if a particle selected was comprised of two nuclei in the original image, or if a particle selected did not appear on the original image. After manual refinement circularity of particles were measured using FIJI (ImageJ) (Schöchlin et al., 2014).

### Pharmacological Inhibitors

Latrunculin B (Sigma, L5288) and Cytochalasin D (Sigma, C8273), Nocodazole (CalBiochem, 31430-18-9) were resuspended using DMSO as a vehicle. Latrunculin B and Cytochalasin D were equilibrated at room temperature for 1 hour prior to use. For experiments inhibitors were diluted to the following final concentrations in 1/9 MR: 1μM Latrunculin B, 10μM cytochalasin D (Lee and Harland, 2007), 150 μM Nocodazole.

### Surface area and volume measurements

IMARIS was utilized to create the 3D renderings and perform the surface area and volume calculations, with a surface area detail level for all treatments of 0.25 μm. Nuclei were excluded if volumes were below 100 μm^3^ or above 1000 μm^3^, as these were determined to be incomplete nuclei or fused nuclei respectively when images were examined.

### CRISPR guide design and injection

CRISPR guides were designed from the V7.1 or V8 gene models on Xenbase and CRISPRscan. Target sites were chosen from UCSC tracks. Guides were chosen using the following criteria: no off targets predicted, a score greater than 50, and in a region in or as close to exon 1 as possible. We generated site specific sg-RNA by ordering a single oligo 5’ - CTAGCTAATACGACTCACTATAGG-(n18) target sequence GTTAGGAGCTAGAAATAG-3’ (Table below). PCR was performed as described in Bhattacharya et al. (Bhattacharya et al., 2015). SgRNA was transcribed using T7 mMachine kit (Ambion). Guides were injected into 1 or 2 cell embryos (dose in table below) with 1.5ng Cas9 (Bhattacharya et al., 2015; Nakayama et al., 2013).

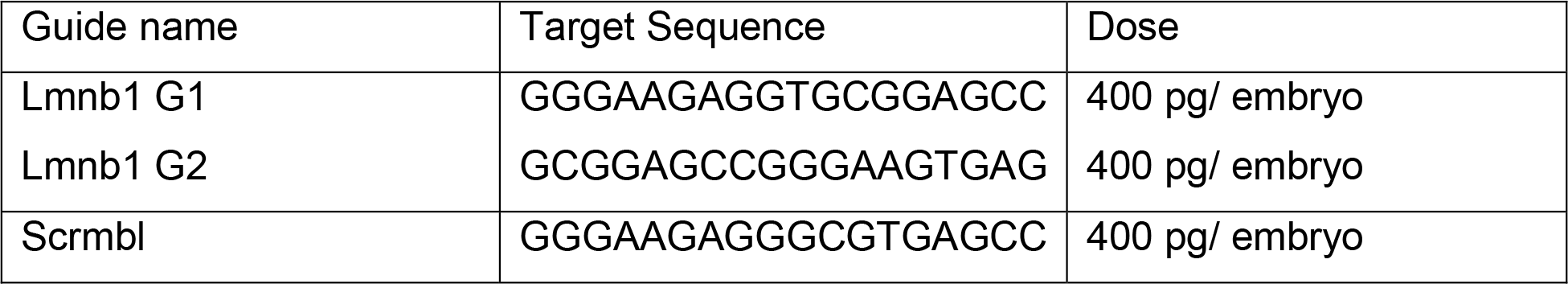

### Dominant negative LmnB1

Lamin B1 dominant negative constructs were constructed following a similar strategy to Schrimer et al. (Schirmer et al., 2001), beginning with *X. laevis* Lamin B1 (Xenbase ORFeome clone XICD00712670) with the following primers: Rod only left primer: GGATCCATGGCCACTGCCACA and right primer: GAATTCCAGTGGCAGAGG.

### High-resolution melt analysis

To extract genomic DNA individual tadpoles were lysed by heating at 95°C in 25mM NaOH and 0.2mM EDTA. Samples were cooled to RT and equal volume of TRIS-HCl 40 mM buffer was added. 2μL of extracted genomic DNA was utilized in PCR reactions containing HRM master mix (GoTaq Flexi buffer (Promega #M8901), dNTPs, MgCl2, DMSO, EvaGreen (Biotium #31000), taq polymerase (Quiagen #201203), nuclease free water). The region of interest was amplified (Left primer GATCTGCAGGAGCTGAATGAC, right primer TGTTCCACGGAGATCTTACTGA) for 35 cycles and melted from 60 - 95°C at 0.1°C increments with EvaGreen fluorescence measured after each temperature change.

### Explants

Dorsal posterior explants were dissected between stage 15-17 as previously described (Tucker and Slack, 2004). Explants were cultured in Danilchik’s for Amy (DFA) buffer without antibiotics. Sibling tadpoles were reared in 1/9^th^ MR as described above.

### Statistical Analysis

R studio was utilized in generating statistics. One-way ANOVA, with Tukey’s post hoc was utilized to calculate P-values for circularity and tail morphometric measurements, for qPCR, nucleoli number, and surface area and volume measurements P-values were calculated with a two-tailed student’s t-test assuming unequal variance.

## Acknowledgements

We thank Ed Parker for assisting with TEM imaging, and the UW vision core facility (NEI P30EY001730). We acknowledge support from the W. M. Keck Center for Advanced Studies in Neural Signaling (NIH S10 OD016240) and the assistance of center manager Dr. Nathaniel Peters. We are grateful to Nathaniel Ng of the Enrique Amaya lab for testing staging conditions for time-lapse analysis and preliminary movies. Alexander Chitsazan helped with training in R and advised on statistical methods. We thank the Molecular and Cell Biology of Xenopus Course at Cold Spring Harbor for embryology and microscopy training, the Xenopus Quantitative Imaging Course at the Marine Biological Laboratories for training in imaging and statistical analysis. We thank Xenbase for curation of genomic and literature information used to generate materials and conduct analysis. We thank Wills lab members, Emily Hatch of FHRC and John Wallingford of UT Austin for comments on the manuscript. Finally, we thank Daniel Levy University of Wyoming, Richard Harland UC Berkley, Cliff Brangwynne Princeton University, and John Wallingford UT Austin for materials.

